# Characterisation of tomato (*Solanum lycopersicon*Mill.) genotypes for morphological and disease resistance traits through cluster analysis

**DOI:** 10.1101/2023.12.13.571404

**Authors:** Uppuluri Tejaswini, Parashivamurthy, R. Siddaraju, K. Vishwanath, T. M. Ramanappa, K. N. Srinivasappa, N. Nagaraju

## Abstract

The characterisation of germplasm is very important for the seed material (breeding) and development of improved varieties. The seed material is collected from the M/s. Noble Seeds Pvt. Ltd., Yelhanka few of them are resistant to ToLCV, where Ty gene background is inserted, five genotypes are non-resistant to ToLCV and one check ArkaVikas (susceptible to ToLCV) conducted the research at Kestur Village Doddballapura during *Kharif*, 2021 and *Kharif*, 2022. The genotypes were morphologically characterised according to DUS guidelines(PPV&FRA). The 20 qualitative and 22 quantitative traits were characterised, out of which 25 traits were observed with variations. Multivariate cluster analysis were assessed for which two clusters (A and B) were formed and for each two, two more sub-clusters (C and D) were observed. The 66.67 % of 30 genotypes (20) are clustered under the Cluster E (15 genotypes), 5 genotypes are considered under the Cluster C, 3 genotypes under the Cluster D and only 2 genotypes are classified under F cluster. The distance in terms of variations is high between the NBLTM-22 and NBLTM-11. And nearing the distance closer and less variations are observed. By this we can say that minimum the distance between the clusters maximum of similarity between the genotypes for the traits.

## Introduction

Tomato (*Solanum lycopersicon* Mill., 2n=2x=24) is the most important Solanaceous vegetable crops grown in every corner of the world. It is mainly used for cooking, processing forms *viz*., salad, puree, paste, ketchup, sauce and soup (Peralta *et al*., 2008). When compared to other fruits and vegetables, tomatoes are low in antioxidant content but routine intake makes a physiologically relevant source of antioxidants and few chemo-protective compounds (Boches *et al*., 2011). To plan for the breeding programmes by breeders there is an importance of characterisation of genotypes and this morphological characterisation is the primary for evaluation of genetic diversity, plays a role in conservation and preservation of resources (Osei *et al*., 2014 and Sacco *et al*., 2015).

The wild tomatoes are important for breeding, as sources of desirable traits for evolutionary studies. The descriptors display a large range of variation, distinct varietal groups and describe phenotypic and morphological diversity with the support of biochemical assessment (Mohan *et al*., 2018 and Pereira-Dias *et al*., 2020). The characterisation consists of characters that are highly heritable which can be distinguished by our naked eye that expresses in all the environment. The evaluation of the phenotypic traits *viz*., plant morphology, leaf characters, fruit morphology, color intensity, firmness etc. are challenging and time-consuming because of the quantitative nature of the traits. Therefore, to identify tomato cultivars with descriptors are even published by International Union for Protection of New Plant Varieties (UPOV, 1992 and Fiorani and Schurr, 2013).

The characterization of genotypes is one of the effective ways to find promising gene sources and utilize them for the creation of improved varieties (Grozeva *et al*., 2020 and Corrado*et al*., 2014). The aim of the current work is to characterize a tomato collection, comprised of 30genotypes using morphological, fruit quality, biochemical and virus resistance traits. Our specific objectives were a) to characterize these genotypes for morphological, fruit quality, biochemical and virus resistance traits; b) identify accessions those have a high yield, enhanced fruit quality and resistance to ToLCV. Selected genotypes with unique and valuable traits could be used in a subsequent breeding program for thedevelopment of tomato varieties with improved fruit quality and high yield.

## Material and methods

a. **Source of seeds –** The seeds are collected from the Noble Seeds Pvt. Ltd., Yelhanka. The seeds of 30 genotypes are differed for resistance against ToLCV of Ty-gene background. 15 lines - Ty-3 gene 05 lines - Without any resistant genes against ToLCV 01 line - Ty-2 gene 03 lines - Ty-6 gene 03 lines - Ty-2 and Ty-3 genes 03 lines - Ty-2, Ty-3 and Ty-5 genes
b. **Experimental design –** The field experiment was carried out at the Kestur village, Doddballapura, Noble Seeds Pvt. Ltd., Yelhanka. Tomato seedlings was transplanted during *Kharif*, 2021 with 25-30 cm, 50 cm and 110 cm plant to plant, row to row, and between row distance, respectively. Application of Fertilizers, irrigation and microclimate were the same for all genotypes. The experiment was conducted in a randomized complete block design (RCBD) with two replications. Every accession was represented by 10 plants in each replicate.
c. **Characterisation -** All genotypes were morphologically for vegetative and reproductive traits, biochemically and ToLCV evaluated for 42 traits by DUS guidelines, PPV & FRA.
d. **Statistical Analysis**-A total of 34 traits including morphological and fruit quality traits were used to establish distinct clusters multivariate analysis of morphological traits using PAST Software.

## Results

### Plant Morphological Traits

The plant morphological traits of 30 genotypes were recorded at different stages of plant growth. The data pertaining to these traits have been presented in table 1-5.

Among 30 genotypes studied, anthocyanin colouration is observed in 13 genotypes and remaining 17 genotypes does not have anthocyanin colouration at seedling stage. The two genotypes have shown determinate type of plant growth habit (NBLTM-12 and NBLTM-23), four genotypes are observed indeterminate growth habit (NBLTM-10, NBLTM-27, NBLTM-29 and NBLTM-30) and remaining 24 genotypes exhibited semi determinate growth habit among genotypes studied. Out of the genotypes studied, 11 genotypes are observed under highly serrated and 19 genotypes are found less serrations on leaf. The 22 genotypes are observed as semi-erect group, seven genotypes are observed with horizontal type and one genotype is observed under semi drooping type of leaf altitude out of 30 genotypes studied. Among genotypes studied, the 22 genotypes are observed for dark green leaves and eight genotypes are noticed green colour. The stem pubescence is present in all 30 genotypes.

The 10 genotypes are observed for weak stem thickness, 14 genotypes considered medium stem thickness and strong stem thickness is noticed in six genotypes. The short intermodal length and medium internodal length is observed in 5 and 23 genotypes respectively. Whereas, longer intermodal length is observed in two genotypes. The 5 genotypes are observed under short stem, medium stem in 23 and 2 genotypes are having long stem at first inflorescence. All 30 genotypes are observed under early flowering habit.

The fourteen genotypes are observed for uniparous type and 16 genotypes are noticed multiparous type inflorescence. Pertaining to the length of inflorescence, short (NBLTM-2) and medium (NBLTM-5) are observed in one genotype each. However, the remaining 28 genotypes are noticed with long length of inflorescence. The few inflorescences are observed in one genotype NBLTM-2 and 14 are noticed medium number and many number of inflorescence are reported in 15 genotypes. The eight genotypes are considered under weak set of fruits on inflorescence, medium fruit set observed in eighteen and the remaining four genotypes are noticed strong fruit set. Weak fruiting is observed in nine genotypes, eighteen genotypes are observed with medium fruiting and three genotypes are noticed strong fruiting. All the 30 genotypes are noticed under yellow colour flower. The non-exerted stigma group is present in all the 30 genotypes. No pubescence on style is found in all the 30 genotypes.

The ten genotypes are observed with smaller calyx, sixteen were medium sized calyx and large calyx was noticed in four genotypes. Anther colour is observed in all the 30 genotypes.

The presence of jointless pedicel is noticed in seven genotypes. Whereas, twenty-three genotypes were absence of jointless pedicel. The six genotypes have been observed for presence of green shoulder and the remaining 24 genotypes are noticed for absence of green shoulder. The absence of fruit depression is noticed in six genotypes, shallow for eleven genotypes, six genotypes are observed for medium fruit depression and for seven genotypes are found for deep fruit depression at the peduncle end. The uniformity of fruits was observed in 26 genotypes and non-uniformity of fruits on plants are noticed in four genotypes.

The flattened shape of fruit is observed in genotype NBLTM-21, the genotypes NBLTM-4 NBLTM-16, NBLTM-25, NBLTM-27, NBLTM-29 and NBLTM-30 were slightly flattened fruit shape. Whereas, circular fruit shape is observed in NBLTM-12, NBLTM-24 and NBLTM-26. Rectangular fruit shape is observed in NBLTM-11 and NBLTM-22. The genotypes NBLTM-3 and NBLTM-17 were cylindrical fruit shape while, NBLTM-1, NBLTM-2, NBLTM-8, NBLTM-9, NBLTM-15 and NBLTM-23 were having heart shaped fruit. The obovoid fruit shape is observed in genotypes of NBLTM-5 NBLTM-7, NBLTM-13, NBLTM-18, NBLTM-19 and NBLTM-28. In NBLTM-14 and NBLTM-20 were observed ovoid fruit shape and pear shaped fruit is found in NBLTM-10.

Intensity of green colour on fruit is light in 16 genotypes, medium intensity was observed in 11 genotypes and in three genotypes dark colour intensity is noticed. Out of 30 genotypes, two genotypes were noticed indented shape, eight were indented flat, seven observed with flat blossom end, seven are found flat-pointed and pointed blossom end is noticed in six genotypes. Pertaining to the colour of fruit at maturity, red colour of fruit was found in twenty-nine genotypes and only one genotype NBLTM-14 was noticed with orange colour. The colour of flesh at maturity, the red colour of flesh was found in twenty-nine genotypes and orange colour flesh was observed in only one genotype NBLTM-14. The fruit weight considered was very small category as the weight of each fruit was less than 100g in all the genotypes. The smaller size at blossom end was observed in one genotype, medium sized blossom end was noticed in 23 genotypes and large sized blossom end was found in five genotypes. The thirteen genotypes were to be considered under the medium sized locules and remaining seventeen genotypes were under large sized locules.

Pertaining to the length of fruit stalk, one genotype was observed with small, eleven genotypes were observed with medium length of fruit stalk and the remaining eighteen genotypes found for the large fruit stalk. The bilocular, trilocular and multilocular were observed in thirteen, eight and nine genotypes respectively. Pertaining to the width of fruits, nine genotypes observed with small, twenty genotypes with medium width and larger fruit width was observed in one genotype NBLTM-25.Pertaining to fruit length, small was observed in sixteen genotypes and medium fruit length was observed in 14 genotypes.

Seven genotypes were observed with medium thickness of pericarp and the remaining 23 genotypes were with thick pericarp. The two genotypes (NBLTM-9 and NBLTM-25) were recorded under medium firmness and remaining all other 28 genotypes were observed under firm fruits. The total soluble solids were found medium in three genotypes, high in eight genotypes and 19 genotypes were observed under very high TSS out of 30 genotypes.All the 30 genotypes were observed under the early maturity.

Response to severity of disease infection among genotypes, eleven were observed under immune, six were resistant, ten were moderately susceptible, two were susceptible and one genotype (NBLTM-30) was highly susceptible.

### Hierarchical Cluster Analysis (HCA)

Cluster analysis (Fig. 1) of all morphological (Qualitative and Quantitative traits) identified into two distinct clusters (A and B). Again cluster A (C and D) and cluster B (E and F) consists of two distinct groups. The 66.67 % of 30 genotypes (20) are clustered under the Cluster E (15 genotypes), 5 genotypes are considered under the Cluster C, 3 genotypes under the Cluster D and only 2 genotypes are classified under F cluster. The distance in terms of variations is high between the NBLTM-22 and NBLTM-11. And nearing the distance closer and less variations are observed. By this we can say that minimum the distance between the clusters maximum of similarity between the genotypes for the traits.

**Fig. 1:**
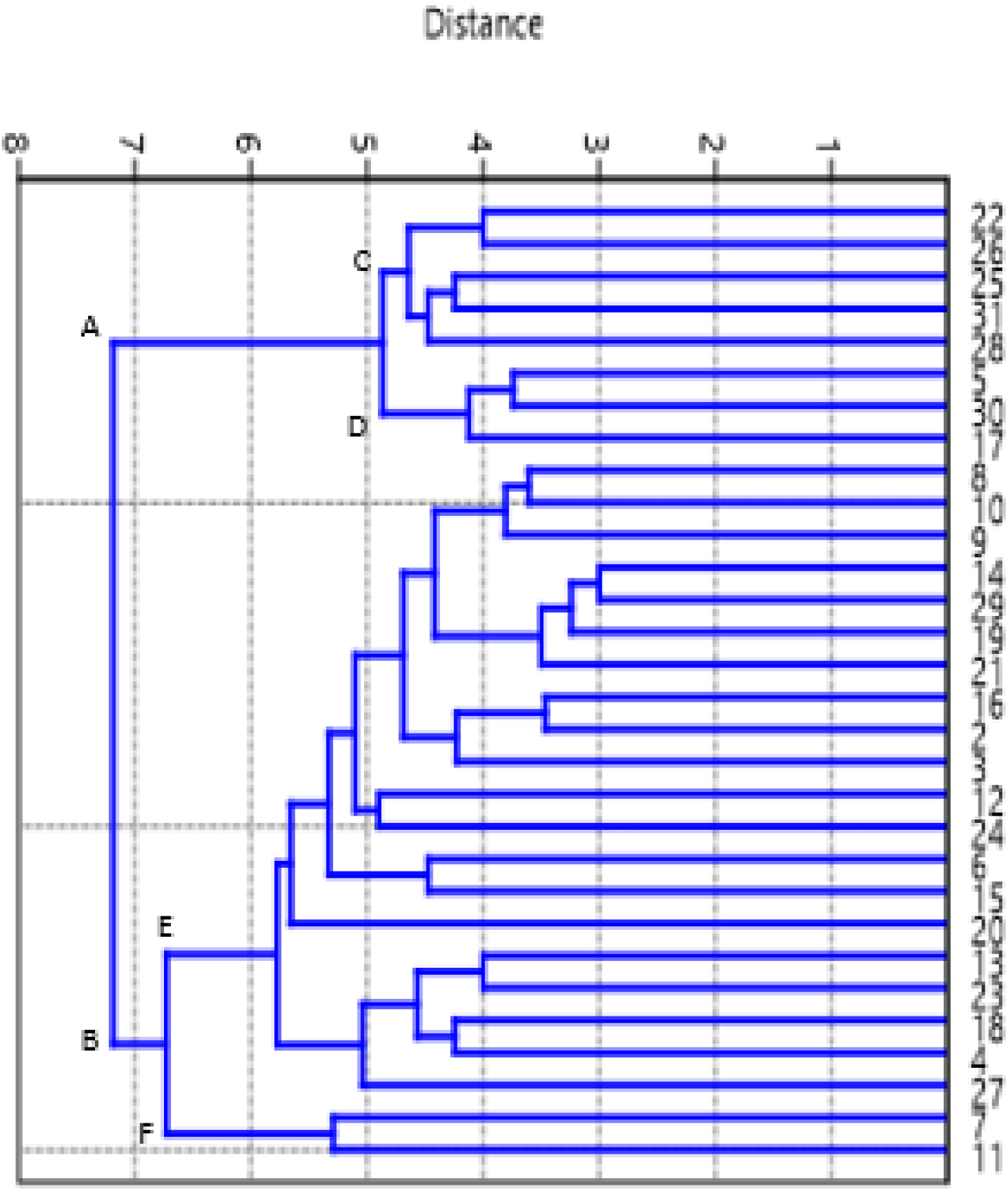
Multivariate cluster analysis of all morphological traits

Cluster C is referred to plant growth habit and leaf altitude. Cluster D (NBLTM-5, NBLTM-30 and NBLTM-17) is confirming for the traits of presence of jointless pedicel, uniformity of the fruits on plant, size at blossom end, size of locules and leaf altitude. Cluster F (NBLTM-7 and NBLTM-11) is similar for the traits *viz*., Anthocyanin colouration, plant growth habit, leaf serration, leaf colour, stem thickness, length of internode and inflorescence, TSS, presence of jointless pedicel and green shoulder. Cluster E consists of many genotypes that is differed by one or the other character but they are closely related. NBLTM-8 and NBLTM-10 is less distant and closely related for all the morphological traits where, NBLTM-8 and NBLTM-27 is very distant and are not closely related for the traits.

These traits are governed by the genotypic inherited traits, but not by the resistance imparted by the gene. The characters are genotype specific and not altered by the resistant gene.

## Discussion

The diversity of a germplasm is critical for the conservation, utilization andvarietal development. The crop improvement program and the varietal development is totarget yield and yield components, but breeding for early maturity, fruit quality and multiple stresses is equally important. To achieve this balance any breeding program needs to have created for future selection (Grozeva*et al*. 2020).

During the course of study 42 traits were reported. Out of these, 20 quantitative and 22 qualitative traits were recorded. Variations were observed in all quantitative characters and few qualitative characters. Only 25 traits showed substantial variations among cultivars and all the quantitative characters found to be useful for characterization of tomato cultivars. Seed, seedling and plant characters are major components of cultivar identification as they provide good data for differentiating characters among genotypes. However, it is difficult to identify genotypes based on single morphological trait. Instead, a set of morphological traits are essential to distinguish the genotypes.

The tomato genotypes differed for developmental, vegetative and fruit traits. The diversity of genotypes is critical for conservation, utilization and development. The prime importance of breeding programme, crop improvement and varietal development that targets to yield and its components that influenced by the morphological traits and innate behaviour of genotypes particularly for breeding early maturity, fruit quality and multiple traits. Conventionally, morphological and agronomic traits have been used in phenotypic evaluation (Grozeva *et al*., 2020; Salim *et al*., 2020 and Nankar *et al*., 2020) and the same sets of traits were also found suitable for detailed accession characterization in this study.

The study reveals the diversity across the evaluated genotypes of tomatoes revealed variations for all the traits of seedling, leaf, fruit characters. The collections have agromorphological variation, which is occurrence of past studies used for agro-morphological traits (Mavromatis *et al*., 2013; Omar *et al*., 2019 and Salim e*t al*., 2020) fruit shape, shape at blossom end (Nankar *et al*., 2020) to characterize the tomato collections. The variability reported for morphometric traits that indicates that producers of tomato prefers peculiar fruit types and the information is used as a base for the development of varieties that has desirable features (Nankar *et al*., 2020). The traits with low variation within the collection are plant growth habit, stem pubescence, days to 50 % flowering, flower colour, style type, style pubescence, anther colour, fruit colour at maturity, fruit weight and days to maturity which are typical for *Solanum lycopersicum*. These findings were on par with the findings of Rajae *et al*. (2018). Similar results with a large variation for some fruit characteristics were also reported in previous tomato studies and these findings were also corroborating with the findings of Sushma *et al*., 2020; Salim *et al*., 2020 and Raj Narayan *et al*., 2020.

The cluster analysis based on 45 agro-morphological traits divided genotypes intotwo distinct groups. Cluster analysis identified accessions those populated in distinct clusters based on their shared similarity and genetic relatedness (Figas et al., 2015 and Nankar et al., 2020). This would likely allow us to use these cluster-specific accessions to breed for specific traits of interest further by developing potential hybrids between these accessions(Grozeva*et al*., 2020). The basis for the establishment of different cultivar groups regardless of their population type and it has been priorly used for cultivar grouping and demonstrated its usefulness in varietal typification across tomato (Díez and Nuez, 2008) and eggplant (Hurtado et al. 2013, 2014). According to fruit morphology, Parisi et al. (2016) distinguish two ‘Sorrento’ morphotypes and Corrado et al. (2014) established that landraces had a genetic structure that is mainly related to fruit type (Willcox*et al*., 2003 and Verheul, 2005).

Our results were similar to previous findings indicating that the fruit size and antioxidants content is negatively linked and small-fruited tomato genotypes possess a higher amount of biochemical compounds. These findings would likely be useful to breeding programs in developing varieties those are enriched with enhanced fruit quality, nutritional components and desirable fruit size.

## Authors contribution

Uppuluri Tejaswini and Parashivamurthy by collaborating with Noble Seeds Pvt. Ltd., Yelhanka, Bangalore framed and designed the project. Parashivamurthy supervised whole experiment. Uppuluri Tejaswini performed the experiments, analyzed the data and wrote the research paper. Parashivamurthy has corrected the final draft.

R Siddaraju, Ramanappa T M, K N Srinivasappa, Vishwanath Koti and Nagaraju N. helped in framing the experiment, supervised, read and approved the final manuscript.

## Declaration or Conflict of interest

All the authors are agreed the manuscript and there is a compatibility between the members.

## Acknowledgement

Uppuluri Tejaswini is thankful to the supervisor for reviewing the careful reading of the manuscript and also providing insightful suggestions. I would like to thank Noble Seeds Pvt. Ltd. for suggestions and I would also thank all the other authors for their continued support to conduct the experiment.

---

| **Genotypes** | **Gene present** | **Genotypes** | **Gene present** |
| --- | --- | --- | --- |
| NBLTM-1 | Ty-3 | NBLTM-16 | No genes |
| NBLTM-2 | Ty-3 | NBLTM-17 | Ty-3 |
| NBLTM-3 | Ty-3 | NBLTM-18 | Ty-3 |
| NBLTM-4 | Ty-3 | NBLTM-19 | Ty-3 |
| NBLTM-5 | Ty-3 | NBLTM-20 | Ty-3 |
| NBLTM-6 | Ty-6 | NBLTM-21 | No genes |
| NBLTM-7 | Ty-3 | NBLTM-22 | No genes |
| NBLTM-8 | Ty-3 | NBLTM-23 | No genes |
| NBLTM-9 | Ty-3 | NBLTM-24 | Ty-6 |
| NBLTM-10 | Ty-3 | NBLTM-25 | No genes |
| NBLTM-11 | Ty-3 | NBLTM-26 | Ty-2 and Ty-3 |
| NBLTM-12 | Ty-2 and Ty-3 | NBLTM-27 | Ty-2 and Ty-6 |
| NBLTM-13 | No genes | NBLTM-28 | Ty-2 |
| NBLTM-14 | Ty-2 and Ty-3 | NBLTM-29 | Ty-2, Ty-3 and Ty-5 |
| NBLTM-15 | Ty-6 | NBLTM-30 | Arka Vikas |

